# RNADecayCafe, a uniformly processed atlas of RNA half-life estimates across multiple human cell lines

**DOI:** 10.1101/2025.08.19.671151

**Authors:** Isaac W. Vock, Dié Tang, Antonio J. Giraldez, Matthew D Simon

## Abstract

RNA expression levels are partially determined by RNA stabilities. Long-lived RNAs accumulate to higher levels than rapidly degraded RNAs when transcribed at the same rate. The extent to which RNA decay contributes to differences in gene expression between genes in the same cell or between different cell types has not been extensively examined. This is in part because reproducible RNA half-life estimates from different biological contexts have not been broadly collected and curated. To address this, we have developed RNAdecayCafe, a database of high-quality RNA half-life estimates in multiple human cell lines. RNAdecayCafe makes use of pulse-label nucleotide recoding RNA-seq data (namely SLAM-seq and TimeLapse-seq), which we found to provide consistent and orthogonally validated half-life estimates. In total, RNAdecayCafe provides half-life estimates from 11 different human cell lines across 15 datasets and a total of 64 samples that were uniformly processed with a set of optimized bioinformatic tools we recently developed (the EZbakR suite). We showcase how this resource reveals the previously underappreciated role of RNA stability in shaping gene expression. RNAdecayCafe will provide a robust database for future studies of RNA decay.

## Introduction

The abundance of a mature RNA in each cell is a function of its synthesis and degradation kinetics (Garneau et al. 2007; Weake and Workman 2010). Regulation of these rates contributes to cell-type expression patterns (Alonso 2012; Fishman et al. 2024). We have an ever-deepening understanding of the regulation of transcription, thanks in large part to copious data from assays probing many aspects of the transcription cycle (ATAC-seq (Buenrostro et al. 2013), ChIP-seq/CUT&RUN/CUT&Tag (Johnson et al. 2007; Skene and Henikoff 2017; Kaya-Okur et al. 2019), PRO-seq (Kwak et al. 2013), etc.). While much is understood about the regulation of RNA degradation (Dowdle and Lykke-Andersen 2025), a relative dearth of data on the kinetics of RNA turnover holds back further progress in the field. For example, while early analyses found the impact of RNA degradation on gene expression to be minimal (Schwanhäusser et al. 2011; Spies et al. 2013; Li et al. 2014), our recent investigation suggested that RNA stability regulation plays a more substantial role in at least one human cell line (Mabin et al. 2025). A well-curated and unified database of RNA turnover measurements from state-of-the-art methods across multiple common human cell lines would provide the field with an important resource to help integrate RNA degradation into our understanding of biological regulation.

Traditionally, RNA stabilities were probed by inhibiting transcription using drugs like Actinomycin D and measuring RNA levels, eventually with RNA sequencing (RNA-seq) (Lam et al. 2001; Yang et al. 2003; Raghavan and Bohjanen 2004; Spies et al. 2013). This method is limited because global transcription inhibition triggers a number of stress responses, which in turn can influence RNA stability thereby confounding these measurements (Jackman et al. 1994; Bensaude 2011; Sun et al. 2012). In addition, the global decrease in RNA levels raises normalization challenges, generally requiring spike-ins to quantify the absolute changes in RNA levels over the transcription inhibition time course (Risso et al. 2014; Chen et al. 2016).

To address some of these challenges, metabolic labeling coupled with biochemical enrichment of labeled RNA was developed (Dölken et al. 2008; Rabani et al. 2011; Tani and Akimitsu 2012). These strategies involve feeding cells with nucleotide analogs that do not globally perturb cellular processes (most widely used are BrU (Tani et al. 2012), 5EU (Jao and Salic 2008), and s^4^U (Cleary et al. 2005)). Labeled RNA can then be biochemically enriched, and the enriched and input samples can be separately sequenced. By tracking the dynamics of the labeled and total RNA, the kinetics of synthesis and degradation can be inferred (Rabani et al. 2011; Neymotin et al. 2014). While this approach eliminates the harsh perturbation of transcription inhibition, it comes with its own challenges. These methods require substantial amounts of input RNA, can introduce biochemical biases during enrichment (especially when using inefficient enrichment strategies), cannot distinguish the desired enriched RNAs from nontrivial levels of contamination, and also are challenging to normalize (Duffy et al. 2015; Lugowski et al. 2018; Duffy et al. 2019; Furlan et al. 2021).

To address these limitations, enrichment-free metabolic labeling approaches were developed (i.e., nucleotide recoding RNA-seq (NR-seq), such as SLAM-seq (Herzog et al. 2017), TimeLapse-seq (Schofield et al. 2018), and TUC-seq (Riml et al. 2017)). These techniques generally combine s^4^U metabolic labeling with nucleotide recoding chemistry to either convert or disrupt the hydrogen bonding pattern of incorporated s^4^U. This leads to apparent T-to-C mutations in the RNA-seq data that indicate sites of s^4^U incorporation, and obviates the need for biochemical enrichment, as reads from labeled and unlabeled RNA can be distinguished bioinformatically. These NR-seq approaches are widely used, provide consistent results when analyzing the same samples (Boileau et al. 2021), and are often considered the state-of-the-art in RNA stability quantification (Duffy et al. 2019; Herzog et al. 2019).

Compiling a robust database of half-lives requires RNA stability assays that yield consistent, high-quality measurements. While NR-seq has several theoretical advantages over other RNA stability assays, we are unaware of any attempts to extensively collect and compare the consistency of half-life estimates from NR-seq and alternative approaches. Half-life estimates from a range of methods were recently compiled from published tables, but NR-seq was almost completely unrepresented in this compendium (Agarwal and Kelley 2022). While that work noted that estimates were plagued by batch effects and poor agreement across methods, it remains to be seen whether independent NR-seq datasets are similarly discordant. In addition, there are two different labeling strategies that can be used in NR-seq experiments: pulse-label (label with s^4^U then extract RNA) and pulse-chase (label with s^4^U, chase with uridine, then extract RNA). RNA half-lives can be estimated with both strategies, but pulse-chases have additional complications (e.g., prolonged s^4^U exposure) that could impact dataset quality (Burger et al. 2013; Neymotin et al. 2014; Altieri and Hertel 2021). We hypothesized that compiling data from NR-seq experiments would thus provide a powerful resource for comparing RNA stability assays and obtaining high-quality half-life estimates.

Towards that end, we compiled nearly all published bulk NR-seq data collected in human cell lines and used our unified and reproducible analysis suite (the EZbakR suite (Vock et al. 2025)) to uniformly reprocess and reanalyze this data. We found that data were most consistent from pulse-label NR-seq experiments. We thus obtained high quality reproducible half-life estimates across 11 human cell lines from the curated pulse-label NR-seq experiments (Supplemental Table 1). To showcase this resource, we examined how RNA stability contributes to both gene-to-gene expression variance in a given cell type as well as gene expression variance across cell types. These analyses revealed that RNA stability regulation impacts steady-state gene expression levels to a much greater extent than previously realized. Together, our database of human RNA half-lives, RNAdecayCafe, represents a powerful resource that promises to deepen our understanding of RNA turnover and how it relates to biological regulation.

## Results

### Datasets and inclusion criteria

We collected, reprocessed, and analyzed publicly available NR-seq data hosted on the SRA at the time of this study (Supplemental Table 1) (Katz et al. 2022). We excluded samples with insufficient s^4^U-induced mutations, resulting in half-life estimates from 136 samples across 28 datasets and 16 cell lines (Methods). Only two of these samples were included in the previous compilation (Agarwal and Kelley 2022).

We first assessed the sample-to-sample correlation in estimates from non-NR-seq data in the Agarwal et al. compendium (Agarwal and Kelley 2022). We hypothesized that cross-dataset estimate concordance may be a useful proxy for dataset quality. Similar to Agarwal et al., we found evidence of frequent batch effects and strong estimate discordance. This included strong disagreements across datasets from the same cell line (Figure 1A). Despite this, we noticed that a subset of samples showed moderate correlation (Pearson > 0.4) across datasets, cell lines, and methods (Figure 1A, rows indicated with purple box). This suggests that obtaining consistent half-life estimates is possible and supports the hypothesis that cross-dataset concordance is an indicator of dataset quality.

**Figure 1.**
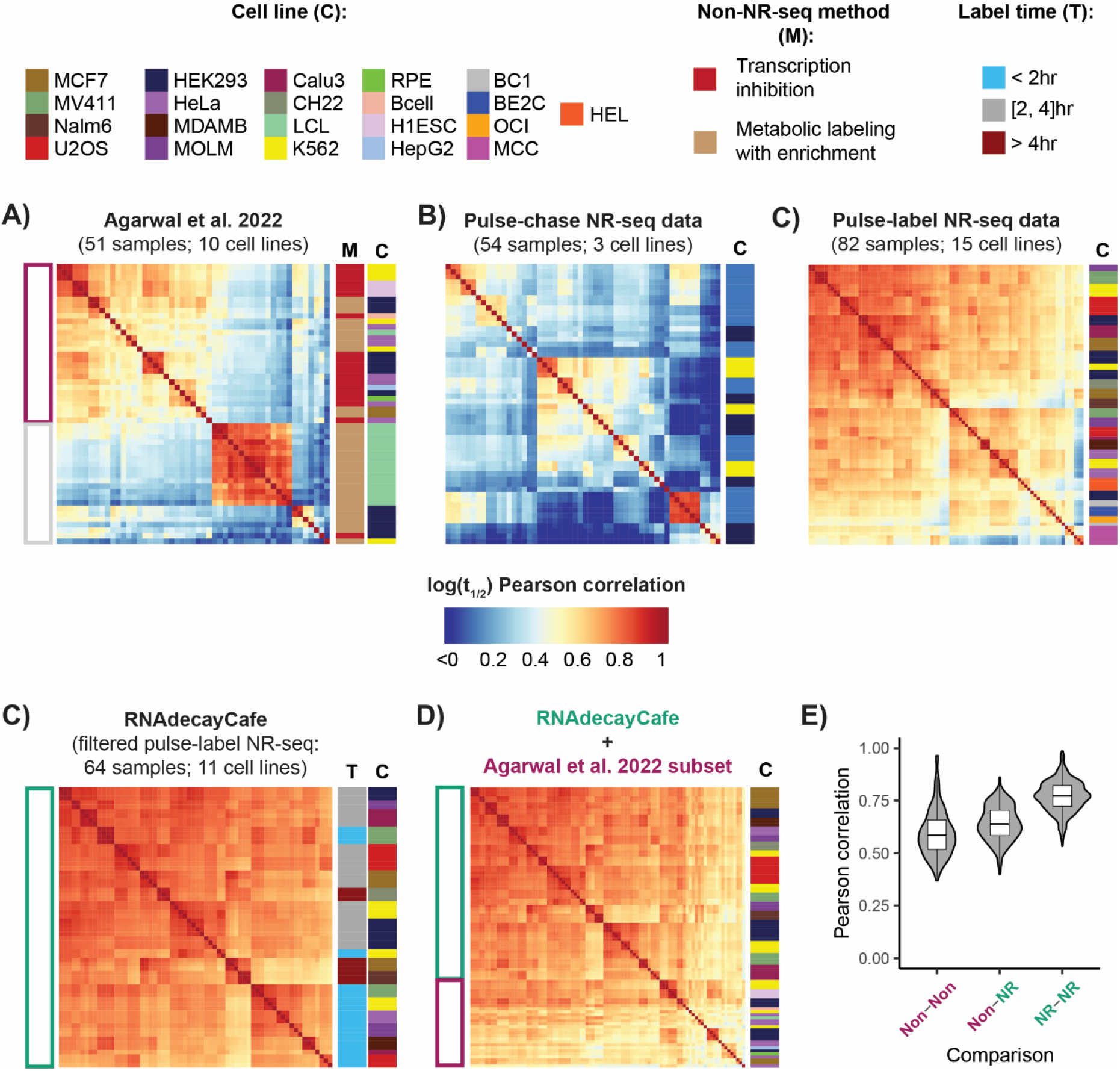
RNAdecayCafe is a high-quality database of human RNA half-lives. **A-D)** Heatmaps of sample-to-sample log(half-life) Pearson correlations (Methods). Annotations on the right side of heatmaps either denote cell line (C), non-NR-seq method class (M), or s^4^U label time used (L) (see legend at top). **A)** Analysis of sample-to-sample log(half-life) correlations adapted from Agarwal et al. 2022. This compendium predominantly includes transcription inhibition RNA-seq and metabolic labeling with biochemical enrichment samples. The three NR-seq samples from that compendium are not included in this heatmap. Moderate-to-high cross-dataset correlation subset denoted with purple box. **B)** Sample-to-sample log(half-life) correlations for all published pulse-chase and pulse-label NR-seq data from unperturbed human cell lines available on the SRA and with detectable s^4^U incorporation (Methods). **C)** Sample-to-sample correlations for RNAdecayCafe datasets, representing a high quality subset of the pulse-label NR-seq data from **B. D)** Comparison of highest concordance non-NR-seq Agarwal et al. half-life estimates to those from RNAdecayCafe. **E)** Comparison of distribution of Pearson correlations between Agarwal et al. non-NR-seq half-life estimates (Non-Non), Agarwal et al. and NR-seq estimates (Non-NR), and NR-seq estimates (NR-NR).

We next assessed the correlation between estimates from NR-seq data. The reprocessed data included NR-seq experiments with two different labeling strategies: pulse-label and pulse-chase (Harrold et al. 1991; Neymotin et al. 2014; Uvarovskii et al. 2019). Half-lives can be estimated with both strategies, but pulse-chases have additional complications (see Discussion) (Nikolov and Dabeva 1985; Burger et al. 2013; Neymotin et al. 2014; Altieri and Hertel 2021). We thus assessed the cross-dataset concordance of pulse-chase and pulse-label NR-seq data separately.

While pulse-chase NR-seq data suffered from even less sample-to-sample concordance than the Agarwal et al. datasets, pulse-label NR-seq data was highly concordant, with almost all samples providing similar relative half-life estimates (Figure 1B and Supplemental Figure 1). This is notable because the diversity of cell types represented in the pulse-label data was higher than that represented in the pulse-chase NR-seq data and the Agarwal et al. dataset (15 cell lines vs. 3 and 10, respectively). After excluding the lowest quality pulse-label NR-seq datasets (Methods), we were left with a highly reproducible set of half-life estimates from 64 samples, 15 different datasets, and 11 human cell lines (Figure 1C). Half-life estimates in this subset are highly consistent across cell lines. The most notable source of variation was the duration of s^4^U labeling (i.e., the label time, Figure 1C row annotation). This was expected, as the label time influences the dynamic range of the half-life estimates (Uvarovskii et al. 2019).

To test if the half-life estimates from filtered pulse-label NR-seq data are unbiased, we compared them with orthogonal estimates from the most cross-dataset-concordant non-NR-seq samples in the Agarwal et al. collection. We found there to be high correlation between the NR-seq and non-NR-seq estimates (Figure 1D). On average, non-NR-seq estimates correlated better with NR-seq estimates than other non-NR-seq estimates (Figure 1E). These observations argue against widespread NR-seq-specific biases as the cause of their high concordance and underscore the power of NR-seq to provide highly reproducible and accurate half-life estimates in a range of human cell lines. Extending on previous comparisons of NR-seq approaches (Boileau et al. 2021; Zimmer et al. 2023), our analysis also reveals the limited impact that NR-seq protocol differences have on the quality of the resulting data, as different recoding chemistries (TimeLapse and SLAM), different sequencing methods (full-length RNA and Quant-seq 3′-end sequencing), and other idiosyncrasies (RNA extraction protocol, s^4^U concentration, sequencing depth, etc.) can yield consistent estimates.

In our database, RNAdecayCafe, we decided to only include estimates from the filtered NR-seq datasets and not the most cross-dataset-concordant non-NR-seq datasets. This is for several reasons: (1) The cross-dataset concordance is higher for filtered pulse-label NR-seq data than for most non-NR-seq data (Figure 1D and E and Supplemental Figure 1); (2) NR-seq data can be processed and analyzed under the same highly optimized bioinformatic and statistical framework (Jürges et al. 2018; Rummel et al. 2023; Vock and Simon 2023; Berg et al. 2024; Müller et al. 2025; Vock et al. 2025); (3) NR-seq is the only class of RNA stability assays with established strategies for bioinformatic detection and correction of common biases (Zimmer et al. 2023; Berg et al. 2024). This facilitates cross dataset integration and allowed us to provide both unnormalized and cross-dataset normalized half-life estimates (Methods). While similar biases are known to inflict transcription inhibition and metabolic labeling with enrichment strategies, correction/normalization is either not possible or requires rarely included controls (e.g., labeled and unlabeled RNA spike-ins). In all, RNAdecayCafe provides the first highly reproducible database of RNA half-lives in a range of human cell lines.

### Using RNAdecayCafe to estimate the contribution of RNA stability to gene expression variance

To examine the potential of RNAdecayCafe to provide novel insights into general features of RNA turnover regulation, we investigated the extent to which RNA stability regulation contributes to different forms of gene expression regulation.

First, we assessed the extent to which RNAdecayCafe validates and generalizes previous reports of RNA turnover regulation. In particular, we investigated recent reports showing that RNAs transcribed from the X-chromosome are on average more stable than their autosomal counterparts in both mouse and human contexts, perhaps owing to differences in the levels of m^6^A in these transcript populations (Rücklé et al. 2023). The original work used SLAM-seq to assess RNA stabilities in mouse ESCs, but RNAdecayCafe provides the opportunity to assess the generality of this conclusion. We first assessed the distribution of half-lives across all samples in RNAdecayCafe and confirmed that on average, X-chromosome-encoded RNAs are more stable (Figure 2A, left). We then stratified this data by cell type and found that this trend was consistently reproduced in all 11 cell types assayed (Figure 2A, right). In addition, the magnitude of the stability difference was similar in all cell types. This highlights the promise of RNAdecayCafe to generalize observations about RNA turnover regulation.

**Figure 2.**
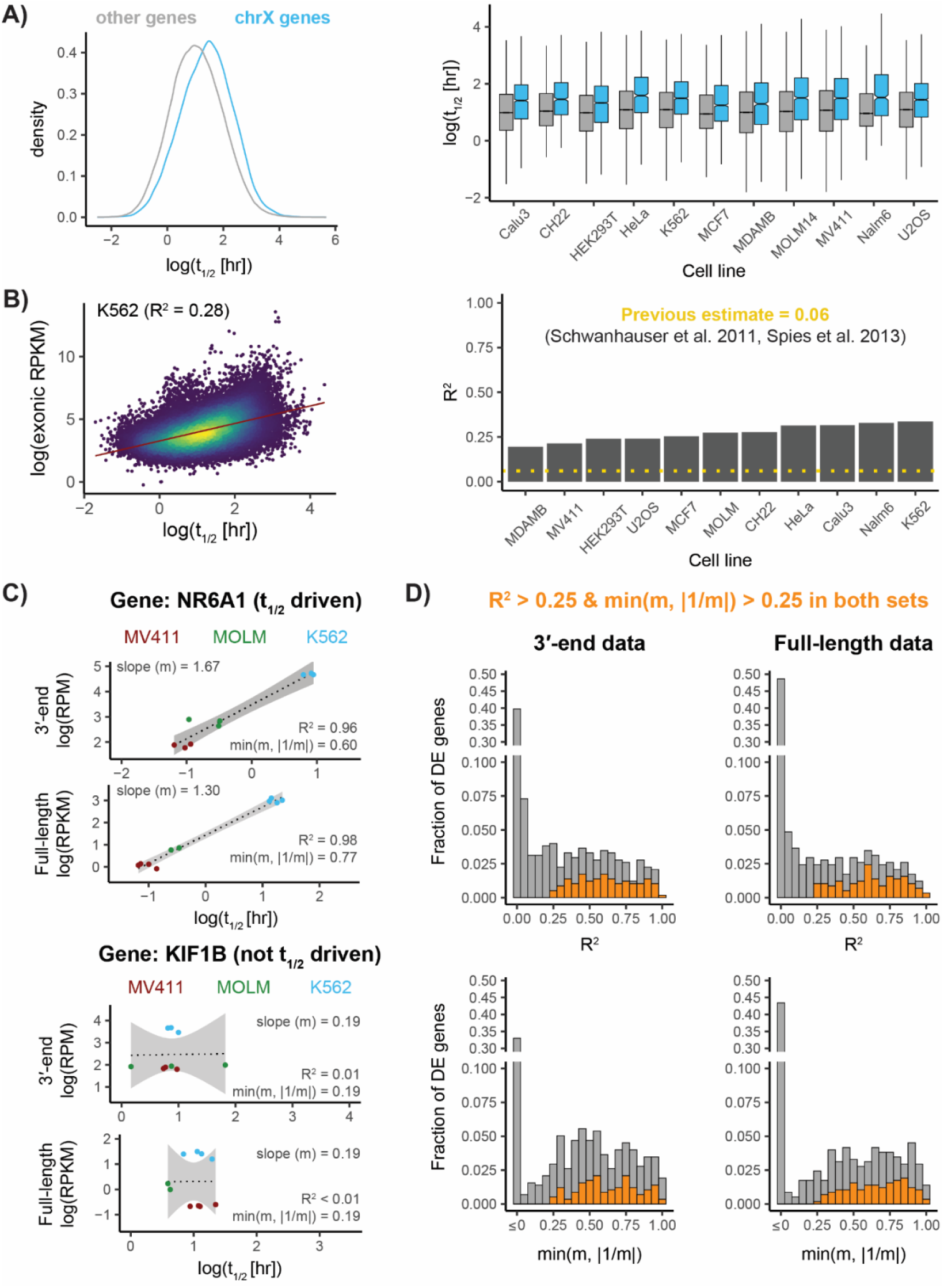
Data from RNAdecayCafe reveals RNA stability regulation. **A)** Analysis of the generality of the previously reported stability of X-chromosome transcripts. Left: compilation of all half-life estimates in RNAdecayCafe. Right: analysis stratified by cell-type. **B)** Analysis of the extent to which gene-to-gene RNA stability differences explain gene-to-gene expression differences. Left: example analysis from K562 data. Right: Coefficient of determinations (R^2^) for all cell lines in RNAdecayCafe. Previous estimates in two different mammalian contexts shown as a yellow dashed line **C)** and **D)** Analysis of the extent to which cell type-expression variance of a gene is explained by that gene’s cell type-stability variance. Analyses constrained to three cell types represented in both full-length and 3′-end sequencing data. **C)** example of a gene (NR6A1), with a strong half-life vs. cell-type expression correlation, and a gene (KIF1B) with weak half-live vs. cell-type expression correlation metrics used in **D** (R^2^ and slope-based metric, see Methods for details) reported. **D)** Distribution of R^2^ and slopes of half-life vs. cell-type expression trends in 3′-end and full-length data. Fraction of genes for which both analyses yielded an R^2^ > 0.25 and slope-metric > 0.25 are noted with orange bars.

Next, we investigated the impact of RNA stability regulation on gene expression. Towards that end, we assessed the fraction of gene-to-gene expression variance explained by gene-to-gene RNA stability variance. Previous estimates ranged between 6-15%, with several independent studies supporting the lower end of this range (Schwanhäusser et al. 2011; Spies et al. 2013; Li et al. 2014). We suspected that noise and biases plaguing older RNA stability assays led to an underestimation of the importance of RNA stability in this setting. We thus assessed the coefficient of determination (R^2^) when correlating log-half lives with log-TPMs in the 11 human cell lines represented in RNAdecayCafe. Analyses in all cell lines yield an R^2^ of around 25%, despite significant differences in sequencing depth and library complexity across samples (Figure 2B and Supplemental Figure 2). We suspect that this still represents an underestimate of the importance of RNA stability. For example, both the half-life and expression estimates have inherent uncertainty arising from finite sequencing depth, s^4^U incorporation rates, etc. In addition, the half-life estimates represent gene-wise averages rather than isoform-specific estimates (Mabin et al. 2025). Despite this, our results demonstrate that previous estimates likely underestimate the impact of RNA stability on gene expression regulation across human cell lines.

Finally, we investigated the extent to which RNA stability explains cell type-specific gene expression patterns. NR-seq datasets in RNAdecayCafe were generated with two different sequencing approaches: full-length RNA sequencing and 3′-end sequencing. We thus focused on the 3 cell types (K562, MOLM, and MV411) which were represented in both full-length (Schofield et al. 2018; Harada et al. 2022; Swartzel et al. 2022; Ietswaart et al. 2024) and 3′-end sequencing datasets (Muhar et al. 2018). We used a conservative ANOVA approach to identify genes which are differentially expressed across these three cell types (Methods). We then assessed the correlation between cell type-specific RNA stability and cell type-specific expression for these genes. Several genes show a striking correlation that suggest their cell type expression trends are almost entirely post-transcriptionally driven, while others show little correlation between their expression and half-life (Figure 2C and Supplemental Figure 3). We quantified the fraction of cell-to-cell expression variability explained by half-life variability (R^2^), and the extent to which half-life variability explained all the expression variability (slope metric, min(slope, |1/slope|); 1 = completely stability driven) for all differentially expressed genes (Figure 2D; Methods). While cell type-specific gene expression is often transcriptionally driven, post-transcriptional regulation also contributes to a significant fraction of the observed expression patterns. In addition, the set of genes with both a post-transcriptional R^2^ > 0.25 and a slope metric > 0.25 was highly concordant between the two analyses (Figure 2D; orange bars). Thus, RNAdecayCafe revealed that RNA stability regulation often contributes to cell type-specific expression patterns.

## Discussion

Here we introduced RNAdecayCafe, a concordant, curated database of RNA half-life estimates from human cell lines. RNAdecayCafe makes use of experimental and bioinformatic innovations, using our recently published EZbakR suite to reprocess published pulse-label NR-seq data, the state-of-the-art method for measuring RNA stabilities transcriptome-wide (Vock et al. 2025). RNAdecayCafe is the first resource of its kind. We thus expect that RNAdecayCafe will contribute to expanding our understanding of RNA stability.

While developing RNAdecayCafe, we found pulse-label NR-seq data to be especially reproducible. While transcription inhibition RNA-seq, metabolic labeling with biochemical enrichment and sequencing, and pulse-chase NR-seq all suffered from notable amounts of dataset-to-dataset discordance, pulse-label NR-seq data consistently provided similar half-life estimates across a wide range of biological contexts (Figure 1 A-C and Supplemental Figure 1). The difference in cross-dataset concordance was particularly striking for pulse-label vs. pulse-chase NR-seq datasets (Figure 1B and Supplemental Figure 1). In pulse-label experiments, cells are treated with s^4^U for the label time, and the RNA is extracted immediately after labeling. In a pulse-chase experiment, cells are first treated with s^4^U for long durations (e.g., 24 hours), and then excess uridine is added as the chase for a controlled chase time. While pulse-chases have been used to great success in the study of biomolecular turnover, we and others hypothesized that they are both unnecessary and suboptimal for metabolic labeling and NR-seq experiments (Harrold et al. 1991; Neymotin et al. 2014). Half-lives can be estimated with both strategies, but pulse-chases have additional complications due to (1) the prolonged exposure of cells to s^4^U (Burger et al. 2013; Altieri and Hertel 2021), (2) nucleotide recycling from degraded RNA during the chase (Harrold et al. 1991; Neymotin et al. 2014), and (3) the need for additional samples (both a pulse and a set of chases) that require additional controls, incur additional costs, and add uncertainty to estimates from pulse-chase experiments. Our results support the hypothesis that these complications can consistently impact the quality of pulse-chase NR-seq data.

The accuracy of estimates from high quality pulse-label NR-seq data is supported by their agreement with estimates from orthogonal approaches (Figure 1D and E) and by their ability to reproduce and generalize a previously reported RNA stability regulation (Figure 2A). We note that even pulse-label NR-seq experiments have potential sources of bias and artifacts, as some datasets were not included because they did not pass quality control, underscoring the need to perform careful QC even with pulse-label NR-seq (Zimmer et al. 2023; Berg et al. 2024). Nonetheless, our analyses suggest that pulse-label NR-seq data provides considerable practical advantages, including higher across-experiment consistency, compared with other high-throughput methods for probing RNA stability.

An important conclusion from our analyses is that RNA stability regulation has a significant and underappreciated impact on gene expression. Gene-to-gene RNA stability variance explains at least one-fifth of the observed gene-to-gene expression variance, and the true fraction is likely higher (Mabin et al. 2025) (Figure 2B). Furthermore, many instances of cell type-specific gene expression are post-transcriptionally driven (Figure 2C, D, and Supplemental Figure 1). While further work will be required to uncover the biochemical drivers of this regulation, our analyses highlight the importance and prevalence of RNA turnover regulation in fine-tuning gene expression.

The high quality half-life estimates across a range of human cell lines provided by RNAdecayCafe opens new avenues for modeling and dissecting the features that influence RNA stability (Agarwal and Shendure 2020; Agarwal and Kelley 2022; Karner et al. 2025). Such efforts promise to further improve our ability to engineer RNAs, explain the impacts of cis-acting genetic variances, and more generally deepen our understanding of RNA biology.

## Methods

### Uniform reprocessing of NR-seq data with fastq2EZbakR

All published NR-seq data from unperturbed human cell lines that was available on the Sequence Read Archive (SRA) was reprocessed with fastq2EZbakR (Katz et al. 2022). Configuration files used for all samples can be found at the Zenodo database hosting all RNAdecayCafe estimates and metadata (link here).

General notes on the preprocessing settings:

1. Multi-mapping reads were excluded from analysis to avoid artifacts
2. In all cases, adapter sequences were trimmed. For 3′-end data, twelve additional nucleotides were trimmed from the 5’end of the reads, as is done by default in the specialized preprocessing pipeline SLAMDUNK (Neumann et al. 2019). For reads from full-length data, three nucleotides were trimmed each end. Quality score end trimming and polyX trimming was also done for all samples.
3. To focus on the half-lives of mature RNAs, only reads aligning exclusively to the union of annotated exons at a gene were used to estimate gene-level half-lives. We used a filtered RefSeq annotation (O’Leary et al. 2016), including only high confidence transcripts identified with sequencing data (i.e., not those that are only computationally predicted).
4. In cases where data was included for no-s^4^U controls, this data was used to call T-to-C SNPs and other mutational anomalies that were not considered when counting T-to-C mutations. This was done with BCFtools mpileup and call (Danecek et al. 2021), with limited downstream filtering to maximize specificity, as implemented in fastq2EZbakR.

### Uniform reanalysis of processed NR-seq data with EZbakR

Processed data from fastq2EZbakR was analyzed with EZbakR. General notes on the analysis settings:

1. If a dataset included no-s^4^U control samples, these were used to estimate a global background mutation rate as well as to correct for dropout using a previously described strategy (Berg et al. 2024). The latter is implemented in EZbakR’s CorrectDropout() function.
2. RNAdecayCafe provides “raw” half-life estimates, as well as those normalized for dropout. Dropout normalization refers to using a proxy for dropout levels to identify the lowest dropout sample (i.e., median estimated turnover kinetics; higher estimates suggest less dropout) and then normalize kinetic parameter estimates relative to this sample using the standard dropout model. Dropout normalization puts half-life estimates from different datasets on the same scale and thus facilitates cross-dataset analyses. Dropout normalized half-life estimates were used in all Figure 2 analyses.

Half-lives were estimated as described previously (Jürges et al. 2018; Schofield et al. 2018; Vock and Simon 2023; Vock et al. 2025) and implemented in EZbakR’s EstimateKinetics function. Briefly:

1. For both pulse-label and pulse-chase data, half-lives were estimated on a per-sample basis. For pulse-labels this means estimating half-lives in each label-fed sample. In pulse-chases this means estimating half-lives in each chase sample.
2. In all cases, half-lives are estimated assuming that the RNA is at steady-state, that RNA is synthesized at a constant rate (k_syn_), and has a single, unchanging rate-constant of degradation (k_deg_).
  a. Given this, the following equations describe a pulse-label experiment with label time tl (R = amount of RNA; θ = fraction of RNA synthesized during the label time, also called the fraction new or the new-to-total ratio):

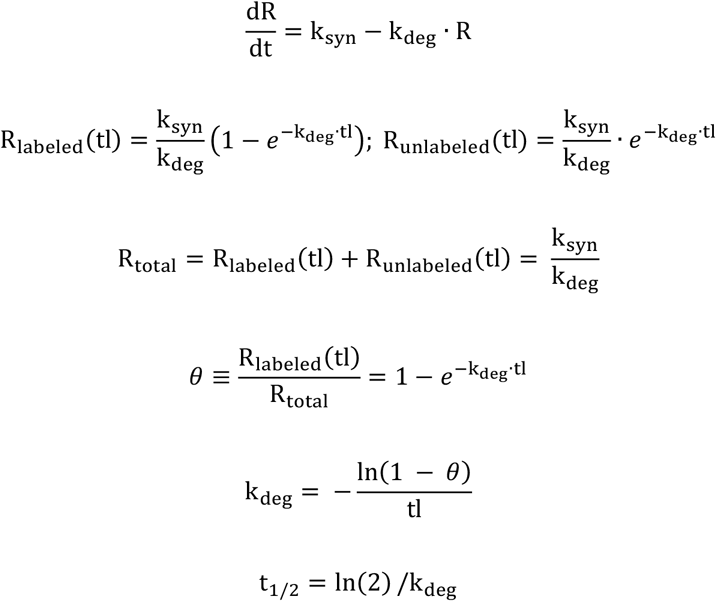
  b. For a pulse-chase experiment with fraction labeled after the pulse θ_p_, fraction labeled after the chase θ_c_, and a chase time of tc, the same differential equation model yields the relationship:

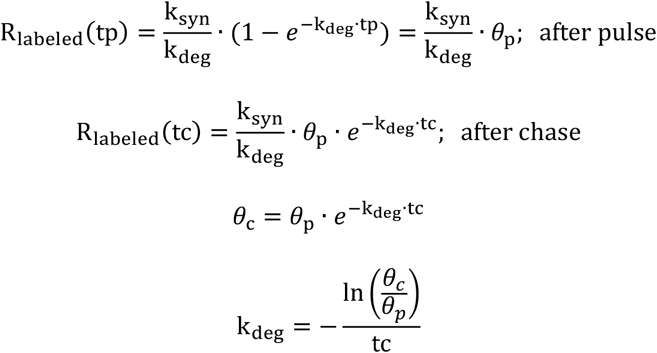
3. For pulse-chases, the average fraction labeled for all replicates of a given pulse time was used to estimate θ_p_. Sometimes, the estimated θ_p_ is less than the estimated θ_c_, in which case the estimated k_deg_ will be negative. In these cases, k_deg_ is estimated assuming that there is one less labeled read in the chase than if the true fraction labeled in the chase were equal to θ_p_ (i.e., that the k_deg_ is near the edge of the dynamic range for that gene).
4. In all cases, EZbakR estimates the fraction of reads from labeled RNA using mixture modeling, as described previously (Jürges et al. 2018; Schofield et al. 2018; Vock and Simon 2023; Vock et al. 2025).

### Sample-to-sample correlation analyses

Sample-to-sample correlation in log(half-life) estimates were computed using R’s cor() function. Pearson correlations were calculated in all cases using the “pairwise.complete.obs” strategy. This means computing Pearson correlations between pairs of samples keeping estimates for genes present in both of the samples. This was important as in some cases (namely the Agarwal et al. dataset), only a small handful of genes are represented in every sample. Rare instances of cross-sample negative correlations were set to 0.

### Quality control filters

For an NR-seq sample to be included in the analyses in Figure 1B, the only filtering done was that the estimated rate of T-to-C mutations in reads from new RNA (referred to as pnew in EZbakR) had to be at least 1%. Values below this typically represent failure of cells to incorporate s^4^U.

For a dataset to be included in RNAdecayCafe, it had to pass the following additional quality control criteria:

1. Limited evidence of dropout. This means that the median, uncorrected half-life estimate should be less than 20 hours, unless there is no-s^4^U data that supports this apparently high stability is not a result of significant amounts of dropout (this caveat did not apply to any samples considered in version 1 of RNAdecayCafe).
2. A pulse-label design had to be used (see Discussion).
3. A label time of at least 1 hour had to be used. Label times shorter than 1 hour are difficult to obtain accurate half-life estimates from as the median human RNA half-life is around 4 hours.
4. Alignment rates had to exceed a minimum threshold of 40%.
5. Good concordance (correlation > 0.75) in RNA-seq read coverage across cell types from similar tissue contexts.

RNAdecayCafe currently includes the following datasets:

1. Schofield et al. 2018 (K562 TimeLapse-seq data; full-length RNA-seq; 4 hour label time) (Schofield et al. 2018).
2. Muhar et al. 2018 (K562, MV411, and MOLM SLAM-seq data; 3′-end sequencing; 1 hour label time) (Muhar et al. 2018).
3. Mowery et al. 2018 (Nalm6 SLAM-seq data; full-length RNA-seq; 5 hour label time) (Mowery et al. 2018).
4. Zuckerman et al. 2020 (MCF7 SLAM-seq data; full-length cytoplasmic RNA-seq. No whole-cell RNA-seq data was collected. Notably, these estimates correlate strongly with those from all whole-cell datasets; 2, 4, and 8 hour label times) (Zuckerman et al. 2020).
5. Luo et al. 2020 (HEK293 TimeLapse-seq data; full-length RNA-seq; 2 hour label time) (Luo et al. 2020).
6. Thiecke et al. 2020 (HeLa SLAM-seq data; 3′-end sequencing; 1 hour label time) (Thiecke et al. 2020).
7. Finkel et al. 2021 (Calu3 SLAM-seq; full-length RNA-seq; 1, 2, 3, and 4 hour label times) (Finkel et al. 2021).
8. Narain et al. 2021 (U2OS SLAM-seq; 3′-end sequencing; 1, 2, and 4 hour label times) (Narain et al. 2021).
9. Sheppard et al. 2021 (CH22 SLAM-seq; 3′-end sequencing; 7 hour label time) (Sheppard et al. 2021).
10. Harada et al. 2022 (MV411 SLAM-seq; full-length RNA-seq; 1 hour label time) (Harada et al. 2022).
11. Swartzel et al. 2022 (MOLM TimeLapse-seq; full-length RNA-seq; 2 hour label time) (Swartzel et al. 2022).
12. Ietswaart et al. 2024 (K562 TimeLapse-seq; full-length RNA-seq; 1 and 2 hour label times; only whole-cell data included) (Ietswaart et al. 2024).
13. Zhou et al. 2024 (HEK293 SLAM-seq; 3′-end sequencing; 3 hour label time) (Zhou et al. 2024).
14. Williams et al. 2025 (MDA-MB-452 SLAM-seq; 3′-end sequencing; 1 hour label time; only whole-cell data included) (Williams et al. 2025).
15. Mabin et al. 2025 (HEK293 TimeLapse-seq; full-length RNA-seq; 2 hour label time) (Mabin et al. 2025).

### Cell type expression vs. t_1/2_ analysis

We restricted cell-type analyses in Figure 2C and D to those genes that showed significant differential expression across cell types. To identify this subset, we performed a conservative ANOVA. ANOVAs typically compare the average within cell line variance to the between cell line variance. We suspected that the assumption of within-group homoskedasticity (same amount of variance for data from each cell line) was likely inaccurate, especially since full-length K562 data came from two separate studies from two separate labs. To account for this, we replaced the average within cell line variance with its max across cell lines. While this will yield a theoretically over-conservative test statistic, we wanted to avoid false positives. Genes with significant cell line-to-cell line expression variance were identified as those with a Benjamini-Hochberg adjusted p-value of less than 0.01 (FDR < 0.01). This left > 2,000 genes for both the 3′-end and full-length RNA datasets.

We used two metrics to assess the extent to which post-transcriptional regulation was responsible for differential expression. One was the t_1/2_ vs. expression R^2^, as used in Figure 2A. While this metric provides rigorous quantification of the fraction of expression variance explained by t_1/2_ variance, it does not capture the fact that the expected relationship between log(expression) and log(t_1/2_) in the case of entirely post-transcriptional regulation is a line of slope 1. Thus, we also devised a slope-based metric to capture the relative amounts of post-transcriptional and transcriptional regulation. For this, we used the minimum of the slope and |1/slope|. This metric has the following properties:

1. Its maximum is 1, when the slope = 1.
2. It gets smaller the further the slope is from 1.
3. It is strictly positive for positive slopes and negative for negative slopes.
4. When the slope is positive, it captures the fraction of the expression range explained by transcriptional regulation. For example, a slope of 0.5 or 2 means that around 50% of the expression range is explained by t_1/2_ regulation. If the slope is 0.5, then there is opposing transcriptional regulation that dampens the effect of t_1/2_ regulation, and if the slope is 2 then there is additional transcriptional regulation that augments the effect of t_1/2_ regulation.

## Data Access

All of the processed data in RNAdecayCafe is documented and provided in a Zenodo repository (link: https://zenodo.org/records/16884513). Half-life estimates can be explored via a shiny app (link: https://isaacvock.shinyapps.io/rnadecaycafe/). Scripts to reproduce figures are hosted on Github (https://github.com/isaacvock/RNAdecayCafe_paper_code).

## Acknowledgements

The authors would like to thank members of the Simon lab, Josh Zimmer, and Florian Erhard for feedback on the manuscript and database. This work was supported by funding from the National Institutes of Health grant T32GM67543-19 (IWV); Alexion Krista Johnson Memorial Fellowship GS102072 (DT); NIH R35GM122580-09 and R01HG013516-01 (AJG); and NIH R01GM137117 (MDS).

IWV and MDS conceived the project with input from DT and AJG. IWV compiled the database, wrote the text, performed all analyses, and made all figures. All authors edited the text. MDS and AJG supervised and secured funding for the project.

